# No evidence for entrainment: endogenous gamma oscillations and rhythmic flicker responses coexist in visual cortex

**DOI:** 10.1101/2020.09.02.279497

**Authors:** Katharina Duecker, Tjerk P. Gutteling, Christoph S. Herrmann, Ole Jensen

## Abstract

Over the past decades, a plethora of studies have linked cortical gamma oscillations (*∼*30-100 Hz) to neuro-computational mechanisms. Their functional relevance, however, is still passionately debated. Here, we asked if endogenous gamma oscillations in the human brain can be entrained by a rhythmic photic drive *>*50 Hz. A noninvasive modulation of endogenous brain rhythms allows conclusions about their causal involvement in neurocognition. To this end, we systematically investigated oscillatory responses to a rapid sinusoidal flicker in the absence and presence of endogenous gamma oscillations using magnetoencephalography (MEG) in combination with a high-frequency projector. The photic drive produced a robust response over visual cortex to stimulation frequencies of up to 80 Hz. Strong, endogenous gamma oscillations were induced using moving grating stimuli as repeatedly done in previous research. When superimposing the flicker and the gratings, there was no evidence for phase or frequency entrainment of the endogenous gamma oscillations by the photic drive. Unexpectedly, we did not observe an amplification of the flicker response around participants’ individual gamma frequencies; rather, the magnitude of the response decreased monotonically with increasing frequency. Source reconstruction suggests that the flicker response and the gamma oscillations were produced by separate, coexistent generators in visual cortex. The presented findings challenge the notion that cortical gamma oscillations can be entrained by rhythmic visual stimulation. Instead, the mechanism generating endogenous gamma oscillations seems to be resilient to external perturbation.

**Significance Statement:** We aimed to investigate to what extent ongoing, high-frequency oscillations in the gamma band (30-100 Hz) in the human brain can be entrained by a visual flicker. Gamma oscillations have long been suggested to coordinate neuronal firing and enable inter-regional communication. Our results demonstrate that rhythmic visual stimulation cannot hijack the dynamics of ongoing gamma oscillations; rather, the flicker response and the endogenous gamma oscillations coexist in different visual areas. Therefore, while a visual flicker evokes a strong neuronal response even at high frequencies in the gamma-band, it does not entrain endogenous gamma oscillations in visual cortex. This has important implications for interpreting studies investigating the causal and neuroprotective effects of rhythmic sensory stimulation in the gamma band.

## 1 Introduction

Cortical gamma oscillations have been repeatedly linked to the formation of neuronal ensembles through synchronization of spiking activity in rodents and primates (e.g. Eckhorn et al., 1988; Gray and Singer, 1989; Engel et al., 1992; Wehr and Laurent, 1996; Brosch et al., 2002), including humans (e.g. Tallon et al., 1995; Müller et al., 1997; Rodriguez et al., 1999; Hoogenboom et al., 2006). Accordingly, they have been ascribed a supporting role for neuronal computations within populations (Singer and Gray, 1995; Singer, 1999; Von der Malsburg, 1999; Engel et al., 2001; Singer, 2009; Nikolić et al., 2013) as well as inter-regional functional connectivity (Bressler, 1990; Varela et al., 2001; Fries et al., 2007). Indeed, numerous studies have been able to link gamma oscillations in the human brain to cognitive processes and perception (see Başar-Eroglu et al., 1996; Herrmann and Mecklinger, 2001; Jensen et al., 2007; Tallon-Baudry, 2009; Uhlhaas et al., 2009, for review), whereas anomalous gamma-band activity has been associated with impaired cognition and awareness, as in e.g. autism spectrum disorder, schizophrenia and Alzheimer’s dementia (see Herrmann and Demiralp, 2005; Uhlhaas and Singer, 2006; Uhlhaas et al., 2009; Traub and Whittington, 2010; Grützner et al., 2013, for review).

In this study, we aimed to entrain, i.e. synchronize, gamma oscillations in the human visual cortex to a rhythmic photic drive at frequencies above 50 Hz. Stimulation at such high frequencies has recently been applied in Rapid Frequency Tagging (RFT) protocols, to investigate spatial attention (Zhigalov et al., 2019) and audiovisual integration in speech (Drijvers et al., 2020), with minimal visibility of the flicker. The ability to non-invasively modulate gamma rhythms would allow to study their causal role in neuronal processing and cognition, as well as their therapeutic potential, as recently proposed by (Iaccarino et al., 2016; Adaikkan et al., 2019).

It is widely accepted that rhythmic inhibition imposed by inhibitory interneurons forms the backbone of neuronal gamma oscillations (Traub et al., 1996; Lozano-Soldevilla et al., 2014, see Bartos et al. 2007; Buzśaki and Wang 2012 for review). Indeed, Cardin et al. (2009) demonstrate evidence for resonance, i.e. a targeted amplification, in the gamma band, in response to optogenetic stimulation of GABAergic interneurons, but not when driving excitatory pyramidal cells (also see Tiesinga, 2012). Here, we ask if a rapid photic flicker can hijack human visual gamma oscillations; a positive outcome would suggest that visual stimulation can modulate pyramidal-inhibitory-network-gamma (PING) activity. To this end, we designed a paradigm that embraces the definition of resonance and entrainment as stated in dynamical systems theory. While neuroscientific studies widely rely on this terminology (e.g. Hutcheon and Yarom, 2000; Schwab et al., 2006; Notbohm et al., 2016; Lakatos et al., 2019), the prerequisites of entrainment are often not sufficiently accounted for, as pointed out by Helfrich et al. (2019). Entrainment requires the presence of a self-sustained oscillator that synchronizes to an external drive (Pikovsky et al., 2003; Thut et al., 2011). This synchronization is reflected by a convergence of the frequency and phase of the endogenous oscillator to the driving force (Pikovsky et al., 2003). Similarly, resonance is reflected by periodic responses to a rhythmic drive and an amplification of individually preferred rhythms, but does not require the presence of self-sustained oscillations per se (Pikovsky et al., 2003; Helfrich et al., 2019). Indeed, studies on photic stimulation at a broad range of frequencies (Herrmann, 2001; Gulbinaite et al., 2019) including the alpha-band (Notbohm et al., 2016) have provided evidence for both resonance and entrainment in the visual system (also see Rager and Singer, 1998, for resonance phenomena in cat visual cortex).

In this study, oscillatory MEG responses to photic stimulation from 52 to 90 Hz were investigated in the presence and absence of visually induced gamma oscillations. In the *flicker* condition, a rhythmic flicker was applied to a circular, invisible patch. In the *flicker&gratings* condition, the flicker was superimposed on moving grating stimuli that have been shown to reliably induce strong, narrow-band gamma oscillations (Hoogenboom et al., 2006, 2010; Van Pelt and Fries, 2013). These oscillations reflect individual neuronal dynamics (Hoogenboom et al., 2006; Van Pelt and Fries, 2013) and have been shown to propagate to downstream areas in the visual hierarchy (Buffalo et al., 2011; Bosman et al., 2012; Bastos et al., 2015; Michalareas et al., 2016). Therefore, we will use the terms *induced* and *endogenous* gamma oscillations interchangeably in the following. We chose moving grating stimuli to elicit narrow-band endogenous gamma oscillations since more complex stimuli induce a broad-band gamma response which might not reflect oscillations (Hermes et al., 2015a,b).

We expected the visual system to resonate to frequencies close the endogenous gamma rhythm elicited by the gratings, as well as a synchronization of the gamma oscillations and the rhythmic flicker. As we will demonstrate, the moving gratings did generate strong endogenous gamma oscillations, and the photic drive did produce robust responses at frequencies up to 80 Hz. However, to our great surprise, there was no evidence that the rhythmic stimulation entrains endogenous gamma oscillations.

## 2 Materials and Methods

### 2.1 Experimental Procedure & Apparatus

The MEG data were recorded using a MEGIN Triux system housed in a magnetically shielded room (MSR; Vacuumschmelze GmbH & co., Hanau, Germany). Neuromagnetic signals were acquired from 204 orthogonal planar gradiometers and 102 magnetometers at 102 sensor positions. Horizontal and vertical EOG, the cardiac ECG signals, stimulus markers as well as luminance changes recorded by a photodiode were acquired together with the neuromagnetic signal. The data were lowpass filtered online at 330 Hz and sampled at 1000 Hz. Structural mag-netic resonance images (MRIs), for later co-registration with the MEG data, were acquired using a 3 Tesla Siemens MAGNETOM Prisma whole-body scanner (Siemens AG, Muenchen, Germany), TE = 2 ms, and TR = 2 s). For two subjects, the T1-weighted images obtained in previous experiments, using a 3 Tesla Philips Achieva Scanner (Philips North America Corporation, Andover, USA), were used (scanned at the former Birmingham University Imaging Centre). Participants were invited to two separate sessions during which the MEG data and the anatomical images were acquired, respectively. Whenever possible, the MEG recording preceded the MRI scan; otherwise, the MEG session was scheduled at least 48 hours after the MRI session to avoid any residual magnetization from the MRI system. Volunteers were requested to remove all metal items (e.g. jewelry) before entering the MSR. To enable later co-registration between MRI and MEG data, four to five head-position-indicator (HPI) coils were attached to the participants’ foreheads. Along with the position of the coils, three fiducial landmarks (nasion, left and right tragus) and over 200 head-shape samples were digitized using a Polhemus Fastrak (Polhemus, Colchester, USA). Following the preparation, the participants were seated in upright position under the dewar, with orientation set to 60*^◦^*. The MEG experiment consisted of fifteen blocks lasting 4 min 30 s each. Participants were offered breaks every *∼*20 min but remained seated. At the beginning of each of these recording blocks, subjects were instructed to sit with the top and backside of their head touching the sensor helmet. The positions of the HPI coils relative to the sensors was gathered at the beginning of each recording block, but not continuously. The MEG experiment lasted *∼*75 min in total.

### 2.2 Rapid photic stimulation

Stimuli were presented using a Propixx lite projector (VPixx Technologies Inc, Saint-Bruno, QC Canada) which allows refresh rates of up to 1440 Hz. To achieve this high-frequency mode, the projector separates the screen (initial resolution: 1920 *×* 1080 pixels) into quadrants and treats them as separate frames, resulting in a display resolution of 960 *×* 540 pixels. The RGB color codes for each quadrant, viz. red, green and blue, are converted to a gray scale, separately for each frame and color, and presented consecutively within one refresh interval. The twelve frames are presented at a refresh rate of 120 Hz, resulting in 12*×*120 Hz = 1440 Hz. This approach allows to drive the luminance of each pixel with high temporal precision, allowing for smooth sinusoidal modulations, reducing unwanted harmonics (see Figure 1C,D). In this study, we applied rapid rhythmic stimulation at frequencies ranging from 52 to 90 Hz in 2 Hz increments.

**Figure 1:**
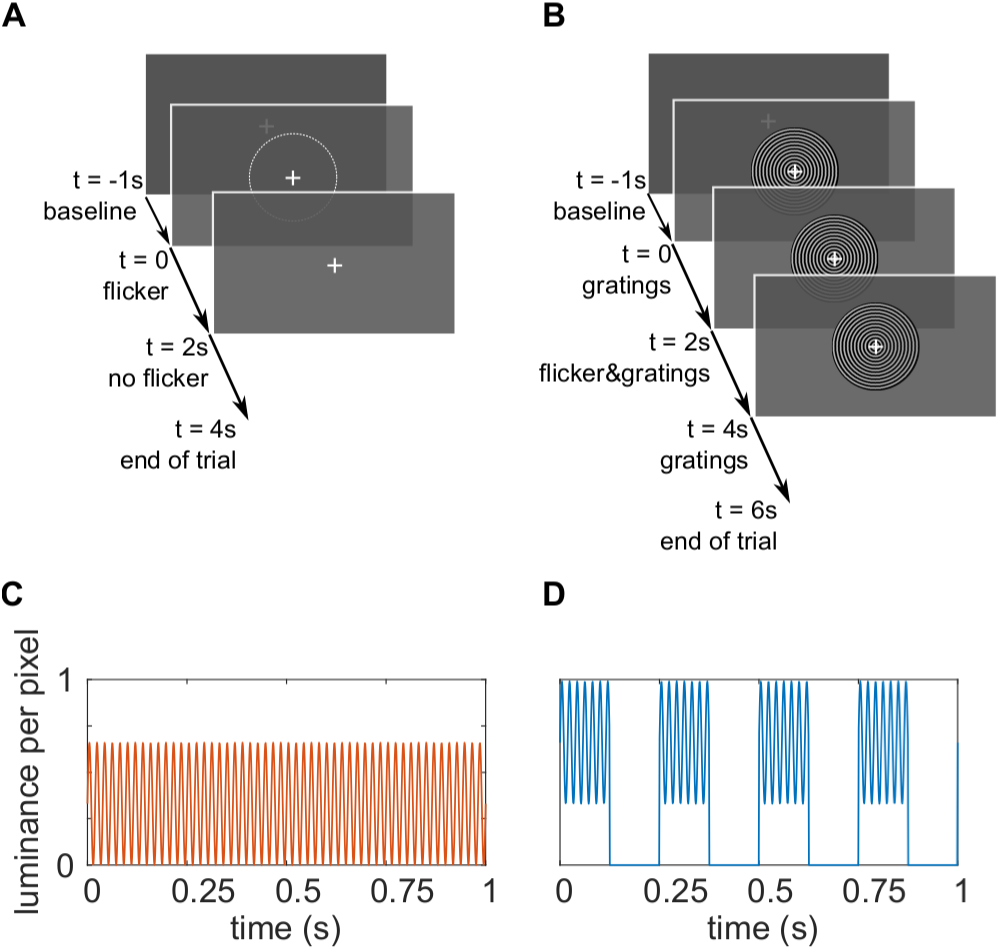
The experimental paradigm. **A** Trials in the *flicker* condition. A 1 s baseline interval with a central fixation cross was followed by a 2 s interval of the rapid flicker applied to a circular patch of size 2.62*^◦^*. The average luminance in the flickering patch was equal to the surrounding gray color, making the photic drive almost unperceivable. The trials ended with 2 s of the fixation cross only. **B** The trials in the *flicker&gratings* condition. The 1 s baseline interval was followed by 2 s of grating stimuli presented centrally on the screen, contracting inwards. Subsequently, the flicker was imposed onto the stimuli for 2 s. The trial ended with a 2 s presentation of the moving gratings without photic stimulation. **C** Sinusoidal luminance change in one pixel induced by the photic drive at 52 Hz in the *flicker* condition. **D** Luminance change in one pixel as a result of the flicker and the gratings moving concentrically with a velocity of 4 cycles/s. To maintain a similar mean luminance between conditions, photic modulation of the invisible patch in **A** ranged from 0 to 66% (mean RGB [84 84 84]), while the light gray rings of the grating, that is 50% of the stimulus’ surface, were flickered between 33 and 99% (mean RGB [168 168 168] per ring).

### 2.3 Experimental Paradigm

Stimuli were created in MATLAB 2017a (The MathWorks, Inc. Natick, MA, USA) and presented using the Psychophysics Toolbox Version 3 (Brainard, 1997). *Conditions* The experiment consisted of two conditions that will be referred to as the *flicker* and the *flicker&gratings* condition, respectively. Each trial began with a one-second interval, in which a central white fixation cross was presented on a dark gray background. In the *flicker* trials, a photic drive in the shape of a circular patch of diameter 2.62*^◦^* was presented for 2 s. Therefore, the patch’s luminance was modulated sinusoidally at frequencies between 52 and 90 Hz (Figure 1A). To minimise the visibility of the flicker, the mean luminance of the patch was matched with the background (33% luminance, RGB [84 84 84]). Frequencies were randomized and balanced across trials. The patch was centered on the fixation cross, such that it was presented both foveally and parafoveally. Each trial ended with a two-second interval in which only the fixation cross was presented. In the *flicker&gratings* condition, the baseline interval was followed by a 2 s presentation of a moving grating stimulus that has been shown to reliably elicit gamma oscillations in visual cortex (e.g. Hoogenboom et al., 2006, 2010; Muthukumaraswamy and Singh, 2013; Tan et al., 2016). The stimulus was the same size as the patch (2.62*^◦^*) and had a spatial frequency of 9.1 rings/*^◦^* (see Figure 1B); the individual rings’ width was 0.11*^◦^*. The rings contracted towards the center of the screen with a velocity of 0.56 *^◦^*/s, i.e. *∼*4.5 cycles/s. In the subsequent 2 s interval, the gratings were flickered at the respective frequencies, by sinusoidally modulating the luminance of the entire stimulus with each screen refresh. The trial concluded with a 2 s interval in which the concentric moving circles remained on screen without photic stimulation. To keep the overall brightness of the stimulation similar between conditions, the luminance of the circular patch in the *flicker* condition ranged from 0 to 66% (of the projector’s maximum), while the brightness of the gratings in the *flicker&gratings* ranged from 33 to 99%. The resulting contrast between the gray and black rings, of 66%, has been previously demonstrated to induce clearly identifiable gamma oscillations (Self et al., 2016). The range of the photic drive, i.e. the difference between peak and trough, estimated based on the projector’s maximum luminance, was 339 lumens. The flicker was replicated in the lower right corner of the screen, to acquire the stimulation signal with a photodiode. The rationale of this design was to investigate if and how the resonance properties of the visual system change when an endogenous gamma oscillator in visual cortex is activated; and whether the flicker response modulates the ongoing oscillatory activity. Studying these two phenomena in the *flicker&gratings* condition required a characterization of both the gamma oscillations and flicker response in isolation. The former was achieved by presenting the gratings without the flicker. To extract the flicker response, we aimed to avoid any gamma-band activity in visual cortex. This was implemented by applying the flicker to a texture-free, invisible patch. Given the filter properties of the visual system (see Cormack, 2005, for review), we were further interested in identifying an upper limit of the frequencies inducing reliable responses. As we expect these results to guide future studies employing the rapid flicker for frequency tagging, we chose an invisible patch to avoid any confounds by response enhancement, e.g. by object-based attention or figure-ground segregation (Self et al., 2016).

#### Task & Time Course

Participants were kept vigilant by performing a simple visual detection task that required them to respond to a 45*^◦^* rotation of the fixation cross at the center of the screen, which occurred once every minute (e.g. Zaehle et al., 2010). Data including the target and/or the responses were discarded and not considered in the analysis. The rotation took place after a trial in the majority, i.e. 60%, of the cases. The remaining 40% of rotations took place at any point during a trial. The experiment was divided into 15 blocks of 4.5 min, resulting in a recording time of 75 min in total. The 40 frequency*×*condition combinations were presented once in each block, in randomized order, resulting in a total of 15 trials per flicker frequency and condition. To minimize the amount of trials rejected by eye-blink artifacts, 3 s breaks, indicated by a motivating catchphrase or happy face on the screen, were incorporated every five trials, i.e. every 25 - 35 seconds. Participants were instructed to utilize these breaks to rest their eyes.

### 2.4 Participants

This project was reviewed and approved by the local Ethics Committee at University of Birmingham, UK. Thirty-one students of the University of Birmingham participated in the experiment. One experimental session was terminated prematurely due to the participant not being cooperative, resulting in a sample of thirty participants (15 female), aged 25.7 *±* 3.4 years. This sample size was decided upon based on a conceptually similar study investigating entrainment of neuronal alpha oscillations by Notbohm et al. (2016). All volunteers declared not to have had a history of neuropsychiatric or psychological disorder, reported to be medication-free and had normal or corrected-to-normal vision. For safety reasons, participants with metal items inside their bodies were excluded at the selection state. Prior to taking part in the study, participants gave informed consent, in accordance with the declaration of Helsinki, to both the MEG recording and the MRI scan and were explicitly apprised of their right to abort the experiment at any point. The reimbursement amounted to £15 per hour. To allow analysis of flicker responses at frequencies with a sufficient distance to the individual gamma frequency (IGF; see 3.1) of the participant, i.e. *±*6 Hz, 8 participants were excluded due to their IGF being below 58 Hz. Thus, the data of 22 participants were included in the following analyses (11 female; mean age 25.7 years).

### 2.5 MEG Data Analysis

Analyses were performed in MATLAB 2017a and 2019b (The MathWorks, Inc. Natick, MA, USA) using the fieldtrip toolbox (Oostenveld et al., 2011).

#### 2.5.1 Sensor Analysis

At the sensor level, the analysis was confined to the planar gradiometer signals, as these provided the best signal-to-noise ratio.

##### MEG preprocessing

Trials containing the target or button presses were excluded. The data were read into MATLAB as 5 s and 7 s trials for the *flicker* and *flicker&gratings* conditions, respectively. Artefactual sensors were identified visually during and after the recordings for each participant, and interpolated with the data of their neighboring sensors (0 to 2 sensors per participant). The individual trials were linearly detrended. Trials containing head movements and/or multiple eye blinks were discarded using a semi-automatic approach. An ICA approach (’runica’ implemented in FieldTrip) was used to project out cardiac signals, eye blinks and eye movement. The sensor positions relative to the HPI coils were loaded in from the data files and averaged for each subject.

##### Time-Frequency Representation of Power

Time-Frequency Representations (TFRs) of power were calculated using a sliding time-window approach (ΔT = 0.5 s; 0.05 s steps). A Hanning taper (0.5 s) was applied prior to the Fourier-transform. This approach induced spectral smoothing of *±*3 Hz. Relative power change in response to the stimulation, i.e. the moving grating and/or the photic drive, was calculated as:

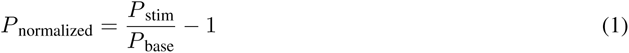

with *P* _stim_ being the power during stimulation and *P* _base_ being the power in the baseline interval. The baseline interval was 0.75 - 0.25 s prior to the onset of the flicker (*flicker* condition) or the moving grating stimulus (*flicker&gratings* condition).

##### Individual Gamma Frequency

The frequency band of the oscillatory activity elicited in response to the moving grating stimulus was identified individually per participant. TFRs of power were calculated for the baseline interval and presentation of the moving grating in the *flicker&gratings* condition and averaged over trials. The results were averaged over the 0.25 - 1.75 s interval, and the frequency bin with the maximum relative power was considered the Individual Gamma Frequency (IGF). For each participant, the 4 to 6 gradiometers with the strongest gamma response to the moving gratings were selected as the Sensors-of-Interest (SOI).

##### Phase-Locking

The average phase-synchrony between the photodiode (recording the visual flicker) and the neuromagnetic signal at the SOI was quantified by the Phase-Locking Value (PLV) (Lachaux et al., 1999; Bastos and Schoffelen, 2016) calculated using a 0.5 s sliding window multiplied with a Hanning taper of equal length. The phases of both signals were calculated from Fourier transformations, applied to the tapered segments. The PLV was computed separately for each *frequency×condition* combination:

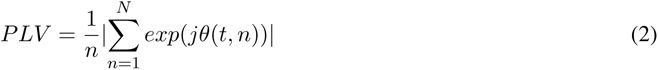

where *θ*(*t, n*) = *φ*_m_(*t, n*) *− φ*_p_(*t, n*) is the phase difference between the MEG (m) and the photodiode (p) signal at time bin *t* in trial *n* (see Lachaux et al., 1999, p.195 and Figure 5 and 9).

**Table 1:** Simple linear regression models: Flicker response magnitude as a function of distance to IGF.

| Model | Estimates |  |  |  |  |
| --- | --- | --- | --- | --- | --- |
| | $\beta_1$ | t | p *** | $R^2$ | F(1,218) |
| *flicker*<sub>plv</sub> | -.01 | -8.07 | $< 2.2e - 16$ | .23 | 65.07 |
| *flicker&gratings*<sub>plv</sub> | -.01 | -7.24 | $< 2.2e - 16$ | .19 | 52.44 |
| *flicker*<sub>pow</sub> | -.07 | -9.01 | $4.80e - 14$ | .27 | 81.14 |
| *flicker&gratings*<sub>pow</sub> | -.16 | -8.95 | $7.51e - 12$ | .27 | 80.13 |

##### Phase difference as a measure of entrainment

Additionally, we investigated changes in phase difference between the photodiode and neuromagnetic signal over time for flicker frequencies of IGF*±*6 Hz, to identify intervals of strong synchrony, so-called *phase plateaus*. MEG and photodiode signals 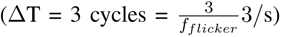 were convolved with a complex Hanning taper using the sliding time window approach. Phase angles were derived from the Fourier transformed time series, unwrapped and subtracted to estimate the phase difference over time for each trial. Plateaus were defined as a constant phase angle (maximum average gradient < 0.01 rad/ms) over the duration of one cycle of the stimulation frequency:

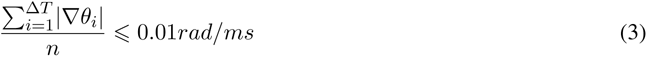

with *∇θ_i_* being the gradient, i.e. slope, of the phase angle between MEG and photodiode signal at a given sample *i*; *n* being the length of the cycle in ms, rounded up to the next integer, e.g. 17 ms for a flicker frequency of 60 Hz. This approach allowed to identify intermittent phase plateaus in each trial. The PLV analysis described above quantifies the phase-similarity of the two signals over trials, and is therefore not feasible to capture brief episodes of synchrony between the MEG signal and the stimulation.

##### Statistical Analysis

Statistical Analysis was performed in RStudio Version 1.2.1355 (RStudio Inc., Northern Ave, Boston, MA; R version 3.6.1., The R Foundation for Statistical Computing).

#### 2.5.2 Source Analysis

##### MRI preprocessing

The raw T1 weighted images were converted from DICOM to NIFTI. The coordinate system of the participants’ individual MRI was aligned to the anatomical landmarks using the head-surface obtained from the MRI and the scalp shapes digitized prior to the recordings. Realignment was done automatically using the Iterative Closest Point (ICP) algorithm (Besl and McKay, 1992) implemented in the FieldTrip toolbox and corrected manually as necessary. The digitized headshape of one participant, for whom there was no anatomical image available, was aligned to a standardized template brain.

##### Linearly Constrained Minimum Variance Beamforming

The neuroanatomical origins of the visually induced gamma oscillations and the response induced by the photic drive condition were estimated using Linearly Constrained Minimum Variance spatial filters (LCMV; Veen et al., 1992), implemented in the Fieldtrip Toolbox (Oostenveld et al., 2011). The MEG forward model was calculated using single-shell head-models, estimated based on the aligned anatomical images, and an equally spaced 4-mm grid, warped into MNI (Montreal Neurologic Institute) space (Nolte 2003, also see Oostenveld et al., 2011; Stenroos et al., 2012); yielding 37,163 dipoles inside the brain. The pre-processed data, epoched in 7 and 5-second trials for the respective conditions, were band-pass filtered at 50 to 92 Hz, by applying second order Butterworth two-pass high- and low-pass filters. To identify the peak locations of the endogenous gamma oscillations and flicker response, respectively, segments of 0.5 s of the baseline interval (0.75 - 0.25 s prior to stimulation) and the stimulation interval (0.75 - 1.25 s after flicker/grating onset) were extracted from the data in both conditions. The peak source of the flicker response to the flickering gratings was isolated based on the 2.75 to 3.25 interval, when the photic drive was superimposed on the gratings, contrasted with the 0.75 to 1.25 interval during which the gratings were presented. For each participant, a common covariance matrix for the 204 planar gradiometers was computed based on the extracted time series and used to estimate the spatial filter coefficients for each dipole location, whereby only the direction with the highest dipole moment was considered. Data in the baseline and stimulation intervals were projected to source space by multiplying each filter coefficient with the sensor time series. Fast Fourier Transforms of the resulting time series, multiplied with a Hanning taper, were computed for each of the 37,163 virtual channels, separately for the baseline and stimulation intervals, and averaged over trials. Relative power change at the IGF and flicker frequencies was computed by applying equation (1) to the Fourier-transformed baseline and stimulation intervals. The source-localized power change values at flicker frequencies up to 78 Hz were averaged to identify a common source for the oscillatory response to the photic drive.

### 2.6 Experimental Design & Statistical Analyses

Using the experimental set up outlined above, this study aimed to explore resonance properties of the visual cortex, reflecting oscillatory dynamics in each participant. Furthermore, we asked if responses to a visual flicker close to and at the IGF are enhanced when the flicker is superimposed on the moving grating stimuli. This would reflect a change in the oscillatory dynamics in presence of the endogenous gamma oscillations. In this context, we hypothesized that these oscillations would synchronize to the flicker. The 40 frequency*×*condition combinations were tested in all participants, i.e. in a within-subject design. Resonance at individually preferred rhythms would be revealed by a high response magnitude to stimulation frequencies in comparison to the surrounding frequencies (Herrmann, 2001; Schwab et al., 2006; Notbohm et al., 2016) (H_1_). A general decrease in response to the flicker as a function of frequency would suggest an absence of such an amplification (H_0_). Entrainment of the ongoing gamma rhythm by the flicker response would result in the peak frequency of the gamma oscillator being synchronized to the stimulation frequency. This is reflected by a reduction in power at the IGF during the application of the flicker to the gratings, at frequencies different from the IGF, compared to the presentation of he gratings alone (H_1_). Statistical analyses were performed in R (R Core Team, 2020, version 3.6.3., using RStudio version 1.2.5033, RStudio Inc., Boston, Massachusetts). The statistical power of the individual tests was evaluated using Bayes Factors, computed using the BayesFactor package in R (Morey and Rouder, 2018). As the identified IGF was found to be higher than the frequency inducing the strongest flicker response in the majority of participants, we quantified their relationship using a simple Binomial test with an a priori defined alpha level of 0.01. The linearity of the flicker response power as a function of flicker frequency, i.e. evidence for the H_0_ as observed in the results reported below, was corroborated using linear regression models implemented in the R base package. Changes in the power at the IGF, with the onset of the flicker in the *flicker&gratings* condition, were examined using a repeated measures ANOVA on the factors time (pre and during flicker) and flicker frequency (above and below IGF), as implemented in package ez in R (Lawrence, 2016). Lastly, we compared the peak sources of the gamma oscillations and flicker responses, identified using LCMV beamforming, in both conditions using dependent sample t-tests. As the direction of the distances was not known a priori, the alpha level was set to 0.025. To reduce the dimensionality of the comparisons, the obtained 3D coordinates were first projected along their first Principal Component (Herrmann et al., 2011). The p-values of the three comparisons were corrected using the Benjamini-Hochberg procedure.

## 3 Results

The aim of the current study was to characterize entrainment and resonance properties in the visual cortex in absence and presence of gamma-band oscillations induced by visual gratings. To this end, we drove the visual cortex with a rapid flicker at frequencies ranging from 52 to 90 Hz, in steps of 2 Hz. The photic drive was applied either to a circular patch (the *flicker* condition, Figure 1A,C) or to the light gray rings of a moving grating stimulus (the *flicker&gratings* condition, Figure 1B,D). We hypothesized that a photic drive in the *flicker&gratings* condition would entrain the grating-induced oscillations. This would be observed as the endogenous gamma oscillation synchronizing with the flicker. Synchronization would be reflected by a constant phase angle between the neuromagnetic signal and the stimulation (’phase entrainment’), as well as a reduction in power at the IGF, indicating a change in the peak frequency of the gamma oscillator towards the flicker frequency (’frequency entrainment’; Pikovsky et al., 2003). Moreover, we expected the presence of the induced gamma oscillator to change the resonance properties (compared to the *flicker* condition), reflected by an amplification of responses to stimulation frequencies equal to the endogenous gamma rhythm. Response magnitudes in the *flicker* condition were expected to reveal resonance properties of the visual system in absence of gamma oscillations, demonstrating favorable stimulation frequencies to be used in future experiments applying Rapid Frequency Tagging (RFT; Zhigalov et al., 2019; Drijvers et al., 2020).

### 3.1 Identifying Individual Gamma Frequencies

The frequency of the endogenous gamma rhythm is known to vary between participants (Hoogenboom et al., 2006, 2010; Muthukumaraswamy et al., 2010; Van Pelt et al., 2012). Therefore, each subject’s Individual Gamma Frequency (IGF) was identified first, based on the 0 - 2 s interval in the *flicker&gratings* condition during which the moving grating stimuli were presented without the visual flicker (Figure 1C). The Time-Frequency Representations (TFRs) of power are depicted in Figure 2A,B for two representative participants. The center column shows the power averaged over time (0.25 - 1.75 s after the stimulus onset to avoid any event-related field confounds) demonstrating distinct peaks at 58 and 74 Hz for these participants. The topographies in the right column depict relative power change at the identified frequencies, focally in sensors over the occipital cortex. For each subject, the 2 - 3 combined planar gradiometers showing maximum relative power change in the gamma band were selected for further analysis (Sensors-of-Interest; SOI) per visual inspection. These sensors strongly overlapped between participants. The data of participants with an IGF closer than 6 Hz to the lowest (52 Hz) drive, i.e. IGF*<*58 Hz, were not considered for further analyses.

**Figure 2:**
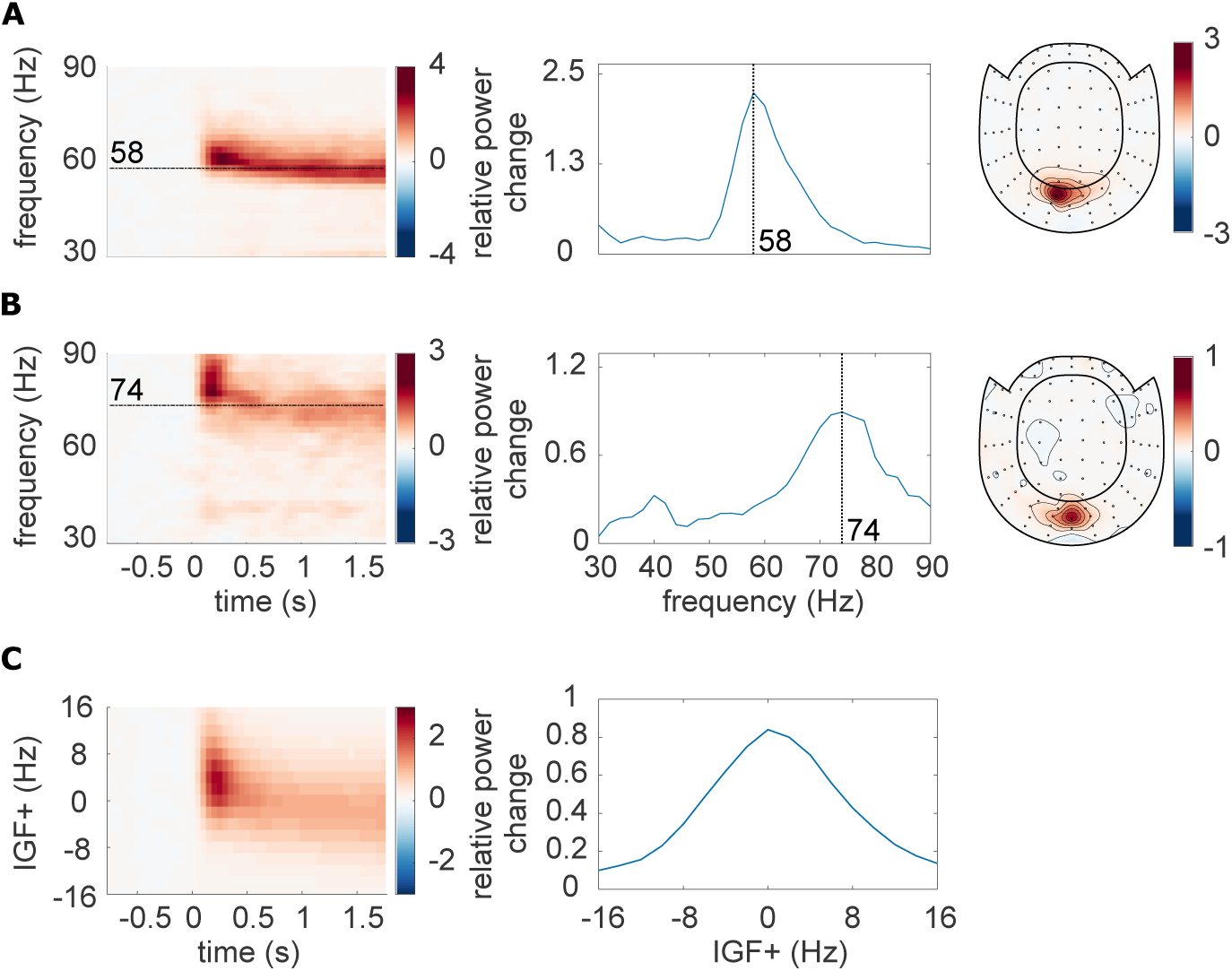
Identification of Individual Gamma Frequencies (IGF) and Sensors-of-Interest (SOI). **A, B** The TFRs of power, power spectra (averaged over 0.25 - 1.75 s) and topographic representations (combined planar gradiometers) of the IGF for two representative participants. The TFRs of power were calculated from the Fourier Transforms using a 500 ms sliding window, resulting in spectral smoothing of ±3 Hz. The IGFs were identified from the spectral peak in 0.25 - 1.75s interval of the TFRs. Identified IGFs are indicated by dashed lines. **C** The grand-average of the power analysis after aligning the individual TFRs and spectra to the IGF (N=22).

Figure 2C depicts the averaged TFRs of power as well as the power spectrum for the remaining subjects (N=22), aligned to each participant’s IGF prior to averaging. The moving grating stimulus induced sustained oscillatory activity constrained to the IGF *±* 8 Hz, with an average relative power change of 80% in the 0.25 - 1.75 s interval compared to baseline. In short, the moving gratings produced robust gamma oscillations observable in the individual participants which reliably allowed us to identify the individual gamma frequencies.

### 3.2 Photic drive induces responses up to 80 Hz

We next set out to quantify the rhythmic response to the flicker as a function of frequency in the *flicker* condition, in which stimulation was applied to an invisible patch. Figure 3 A and B, left panel, depicts the overlaid power spectra for the different stimulation frequencies in two representative participants (the same as in Figure 2). The spectra were estimated by averaging the TFRs of power in the 0.25 - 1.75s interval after flicker onset. Due to the overlap of the sensors detecting the gamma oscillations and photic drive response (compare Figure 2 and 3 right columns) the same SOI were used as in the *flicker&gratings* condition. Both individuals showed strong responses at the respective stimulation frequencies, with a maximum relative power change of 200% and 500% in subject A and B, respectively. The identified IGFs (indicated by vertical dashed lines) were higher than the frequencies inducing the strongest flicker response in 20 out of 22 participants (exact Binomial Test against *H*_0_: *p* = 0.00012, probability of successes (IGF>flicker freq) 0.91*, BayesFactorBF*_10_ = 309.3). When averaged over all participants, the magnitude of the flicker response decreased systematically with frequency (Figure 3C).

**Figure 3:**
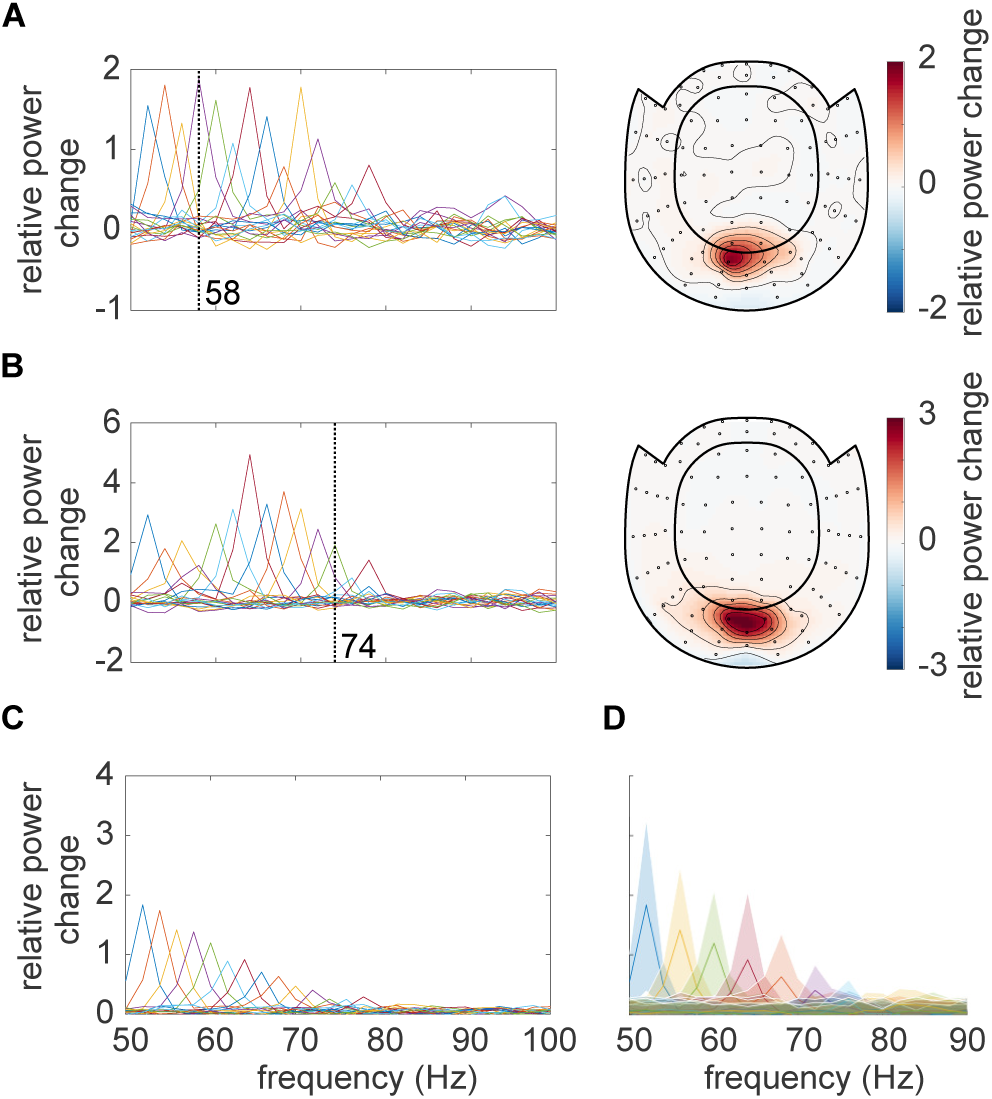
**A,B** The response to the photic drive in the *flicker* condition and the corresponding topographies for two representative subjects. Spectra were estimated from the TFRs of power averaged in the 0.25 - 1.75 s interval. Dashed vertical lines indicate the participants’ IGF. The topographies (combined planar gradiometers) demonstrate a strong overlap with the ones in Figure 2. **C** Grandaverage of the responses to the photic drive for each flicker frequency. On average, the magnitude of the flicker response decreases with increasing frequency, and is identifiable for stimulation below 80 Hz. **D** Grandaverage flicker responses for frequencies from 52 to 90 Hz in steps of 4 Hz. The shaded areas, illustrating the standard deviation, indicate a substantial inter-subject variability.

Figure 4A displays the power spectra in the *flicker* condition, estimated from the TFRs as explained above, averaged over all participants, as a function of stimulation frequency. These are equivalent to 3C. Diagonal values indicate the magnitude of the oscillatory responses (relative to baseline) at the stimulation frequencies, reaching values of up to 300% and decreasing monotonically with frequency. This confirms an upper limit for the stimulation of around 80 Hz. Off-diagonal values indicate oscillatory activity at frequencies different from the stimulation frequency. Figure 4B shows the same spectra after aligning to the individual IGFs, prior to averaging. Figure 4C and D display the spectra in the *flicker&gratings* condition (averaged in the 2.25 - 3.75s interval), during which the photic drive was applied to the moving grating stimulus (see Figure 1B). The induced gamma band activity can be observed as the horizontal light red band at *∼*60 Hz. When aligning the spectra to the IGF (Figure 4D), we observe a decrease in the flicker response but no evidence for an amplification at or close to the IGF.

**Figure 4:**
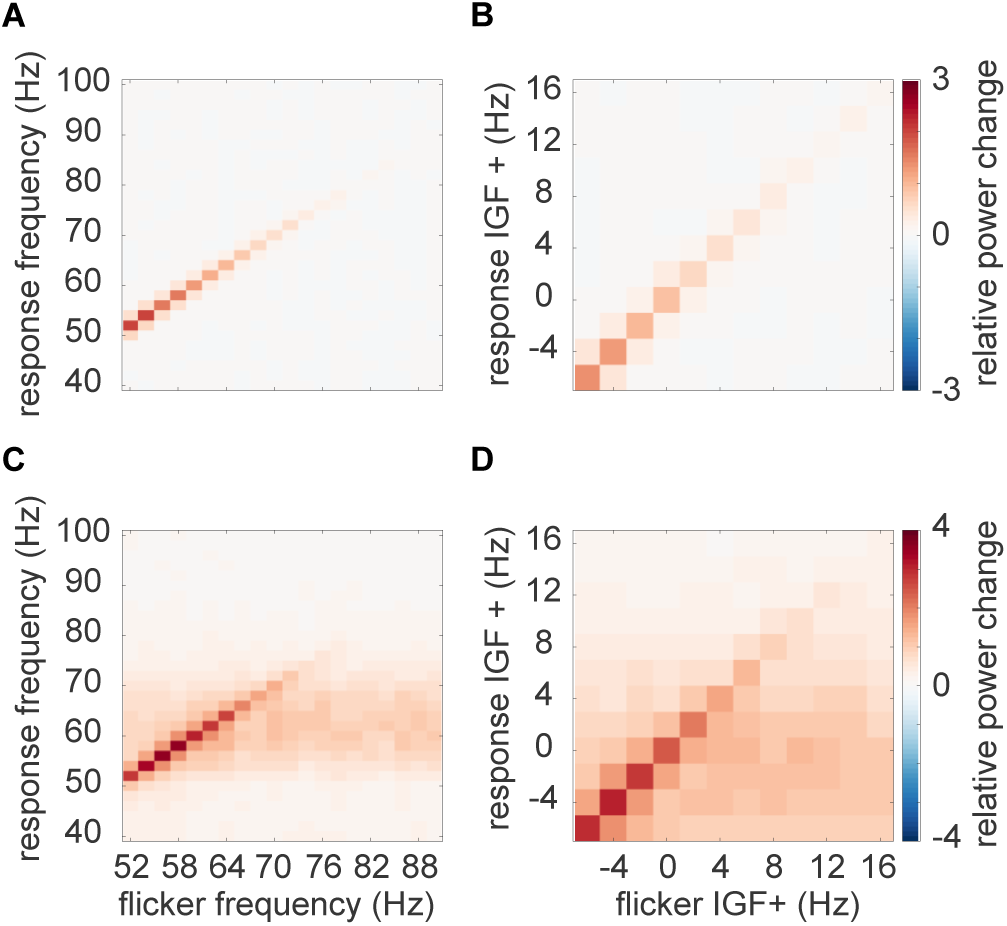
Average relative power change to the photic drive (y-axis) with respect to the driving frequencies (x-axis). **A** The *flicker* condition. Note that the power changes mirror Figure 3C. Power decreases with increasing frequency, from a relative change of 3 at 52 HZ to .5 at 80 Hz. **B** The *flicker* condition after the spectra were aligned to the IGF. **C** The *flicker&gratings* condition. All spectra demonstrate both the flicker response and induced gamma oscillation (observed as the light red horizontal band). Again, the amplitude of the rhythmic stimulation response appears to decrease with increasing frequency. **D** The spectra for the *flicker&gratings* condition now aligned to the IGF. There is no indication that the rhythmic flicker captures the endogenous gamma oscillations.

### 3.3 Magnitude of flicker response decreases as a function of frequency

The averaged TFRs of power in Figure 4 point to an approximately linear decrease in power of the flicker response with increasing frequency. Literature on neural resonance and entrainment, however, suggests the existence of a preferred rhythm at which oscillatory responses are amplified (Hutcheon and Yarom, 2000; Herrmann, 2001; Pikovsky et al., 2003; Notbohm et al., 2016; Gulbinaite et al., 2019). As argued in Pikovsky et al. (2003) phase-locking between the driving signal and the self-sustained oscillator is the most appropriate metric to investigate entrainment. Figure 5A,B depicts the phase-locking value (PLV) between the photodiode and the MEG signal at the SOI (planar gradiometers, not combined). This measure reveals a systematic decrease in phase-locking with increasing flicker frequency for both the *flicker* (orange) and *flicker&gratings* (blue) condition (A). The observed relationship is preserved when aligning the frequencies to the IGF (B, also see Table 1). Note the absence of increased phase-locking at the IGF. The magnitude of the flicker response, quantified by power change compared to baseline, as a function of frequency, is demonstrated in Figure 5C-F and depicts a similar relationship to the one observed for the PLV. The *flicker* condition (C, orange line) revealed a systematic decrease with frequency, whereas the *flicker&gratings* condition did show a peak at 56 Hz. However, this observed increase appeared to be caused by considerable variance between the power estimates of the individual participants (see Figure 5E, each line graph depicts power estimates per individual participant). We again aligned the spectra to the IGF before computing the grand-average (Figure 5D). The absence of a peak at 0 Hz suggests no evidence for resonance at the IGF, confirming the peak at 56 Hz in C to be the result of inter-subject variability. Indeed, simple linear regression models, fit individually to PLV and power as a function of frequency aligned to the IGF, separately for each condition, explain a considerable amount of the variance (see Table 1 and dotted lines in Figure 5). We then identified the individual peak frequencies, eliciting the strongest response to the flicker in the *flicker&gratings* condition 5E, and related those to the IGF, as seen in Figure 5F. As observed in the *flicker* condition, the frequency inducing the strongest response to the flicker was lower than the IGF in the majority of participants, i.e. 19 out of 22 (exact Binomial Test against *H*_0_ : *p* = 0.0008, Bayes Factor *BF*_10_ = 67.5).

**Figure 5:**
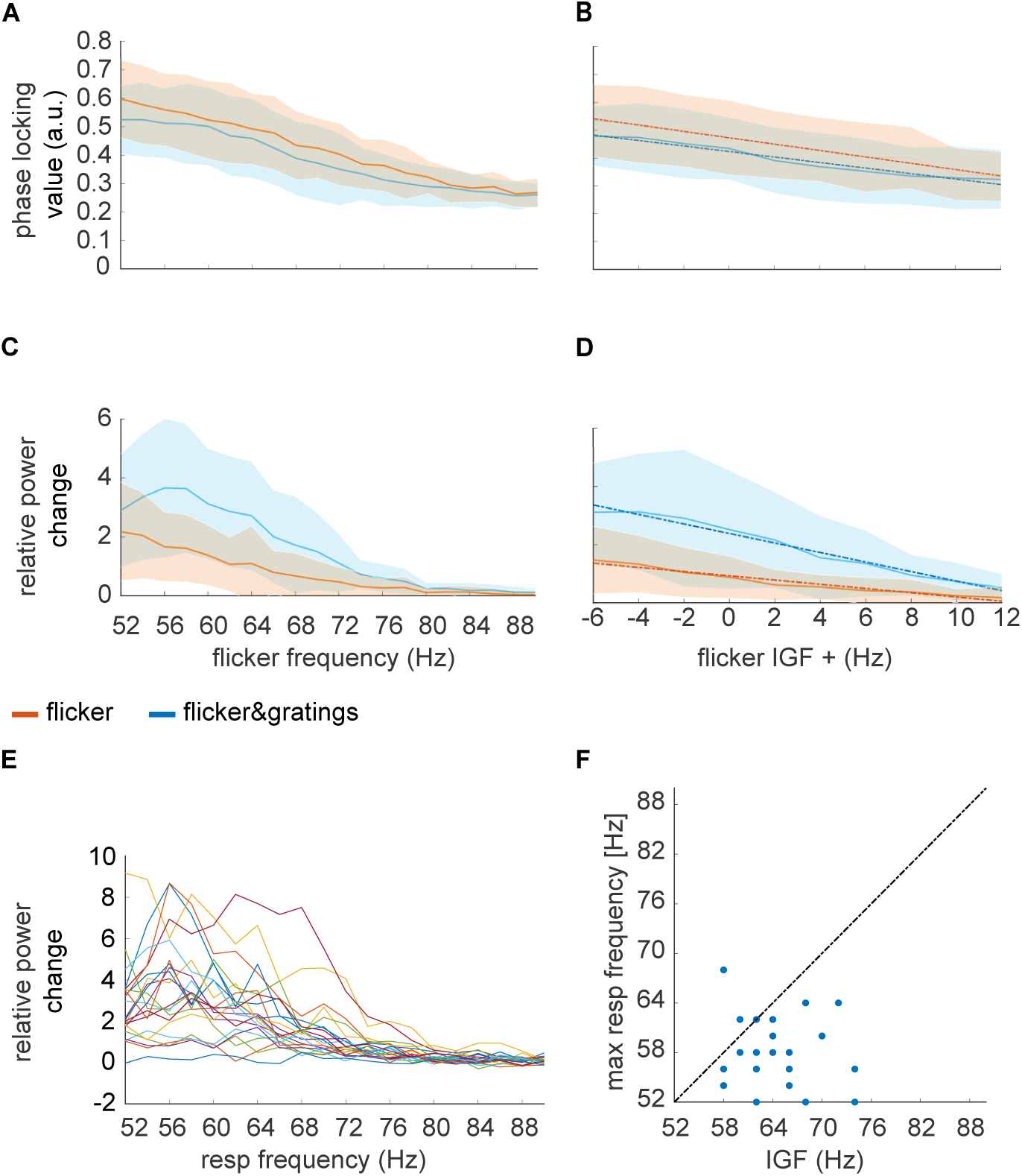
Magnitude of the flicker response as a function of frequency in the *flicker* (orange) and *flicker&gratings* (blue) condition. Shaded areas indicate the standard deviation. **A** The phase-locking values between the photodiode and the MEG signal over the SOIs as a function of driving frequency. **B** The phase-locking values between the photodiode and the MEG signals as a function of frequency after the spectra were aligned to the IGF. Again, the phase-locking decreases with increasing frequency (see Table 1 for a statistical quantification of the simple linear regression models). **C** Relative power change with respect to baseline as a function of frequency. Generally, the power decreased with frequency, however, in the *flicker&gratings* condition there is an apparent peak at ~56 Hz. The shaded areas (standard deviation) indicate considerable variance between participants. **D** Relative power change as a function of frequency after the individual spectra were aligned in frequency according to the IGF, demonstrating that responses to a photic drive at the IGF are not amplified. **E** Relative power change as a function of frequency for each individual subject (N = 22), indicates that the peak at ~56 Hz in **C** is driven by comparably high power in that frequency range in just a few individuals. **F** Flicker frequency inducing highest power values versus IGF, demonstrating the IGF to be higher than the frequency inducing maximum power change in the majority of participants.

### 3.4 Gamma oscillations and flicker response coexist

We initially hypothesized that entrainment of the gamma oscillations in the *flicker&gratings* condition would result in the photic drive capturing the oscillatory dynamics when the driving frequency was close to endogenous gamma oscillations. Figure 6 depicts the TFRs of power relative to a 0.5 s baseline, for one representative subject (also shown in Figure 2 and 3A). The averaged trials for a photic drive at 52 Hz are shown in Figure 6A and separately for each flicker frequency in Figure 6B (Figure created using function by Kumpulainen, 2020). The IGF (58 Hz for this subject) and the respective stimulation frequencies are indicated by dashed lines. The endogenous gamma oscillations, induced by the moving grating stimulus, are observed as the sustained power increase from 0 - 6 s whereas the flicker response is demonstrated by a power increase at 2 - 4 s. The plots reveal that gamma oscillations persist at the IGF and coexist with the response to the photic drive, which is particularly apparent for stimulation at 52 Hz (Figure 6 A). Furthermore, the power increase at the flicker frequency does not appear to outlast termination of the drive at t = 4 s. In the subsequent step, we frequency-aligned the TFRs of power according to the IGF before averaging over participants. Again, the analyses were constrained to individuals with an IGF above 56 Hz (N = 22).

**Figure 6:**
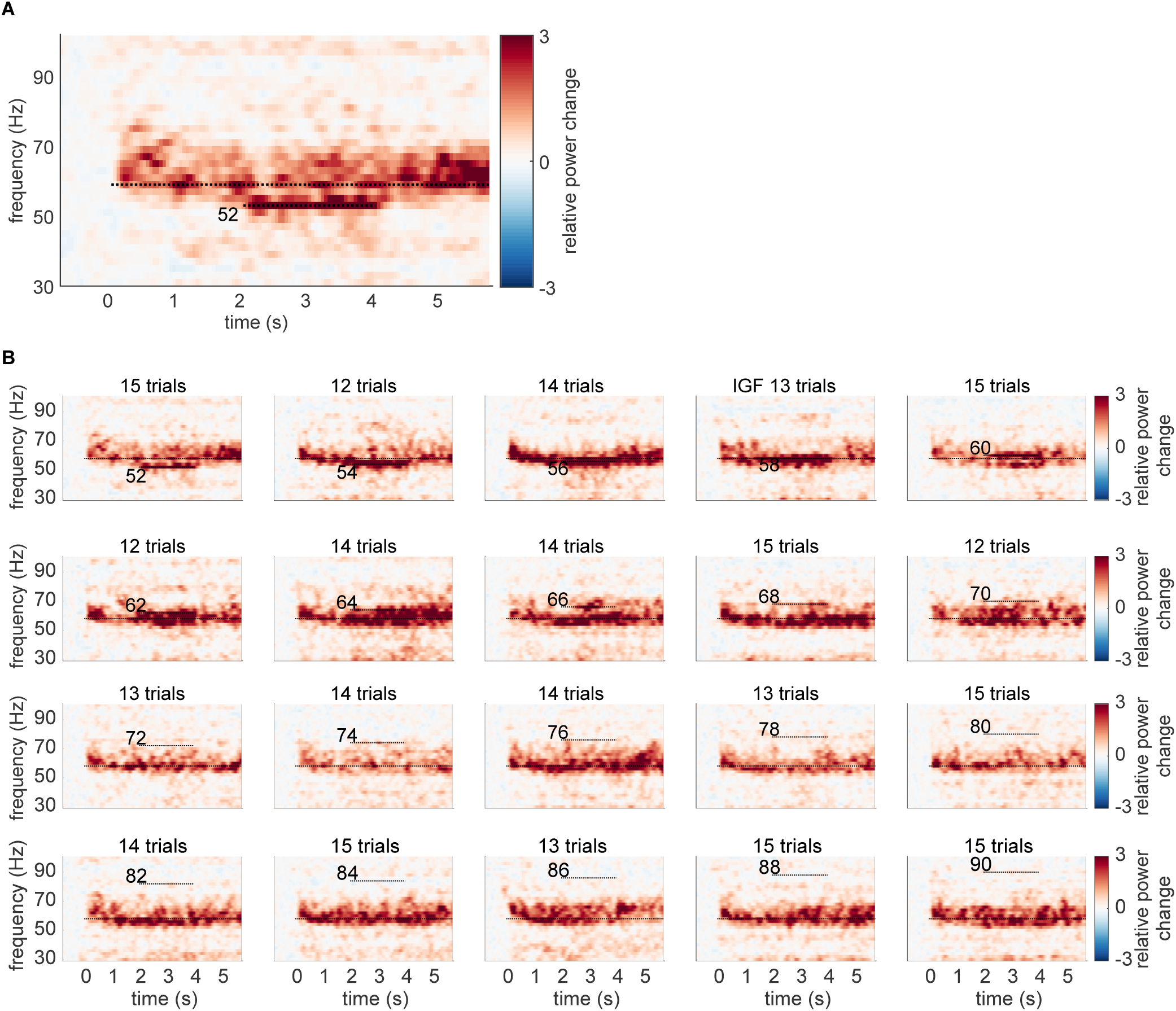
The time-frequency representations (TFRs) of power for one representative subject, showing relative power change averaged over trials and SOIs in the *flicker&gratings* condition. **A** Photic drive at 52 Hz. The moving grating stimuli were presented for 0 - 6 s, with the flicker superimposed from 2 to 4 s. Sustained gamma-band activity is clearly observable throughout the presentation of the stimuli, with a power increase of 300% relative to baseline. Additionally, the rhythmic stimulation elicited a response at 52 Hz, which seems to coexist with the gamma oscillations, indicating that the photic drive is unable to capture the dynamics of the gamma oscillation. **B** The plots for the frequencies from 52 to 90 Hz. Stimulation frequencies and IGF (here 58 Hz) are indicated by horizontal dashed lines. The flicker induced responses up to 66 Hz in this participant. Gamma oscillations persist in presence of flicker responses, suggesting that they coexist.

The group averaged, aligned TFRs are shown in Figure 7 for frequencies ranging from IGF-6 Hz to IGF+16 Hz. The endogenous gamma oscillations are observed as the power increase extending from 0 - 6 s, and the flicker response as the power change in the 2 - 4 s interval marked by dashed lines, respectively. The photic stimulation induces a reliable response that decreases toward 12 Hz above the IGF. Despite the representation of the gamma oscillations being smoothed due to inter-individual differences, the averaged aligned TFRs of power support the observations in the single subject data: both the gamma oscillations and flicker response coexist in the 2 - 4 s interval. Furthermore, there is no indication of the gamma power being reduced during RFT at frequencies close to, but different from, the IGF.

**Figure 7:**
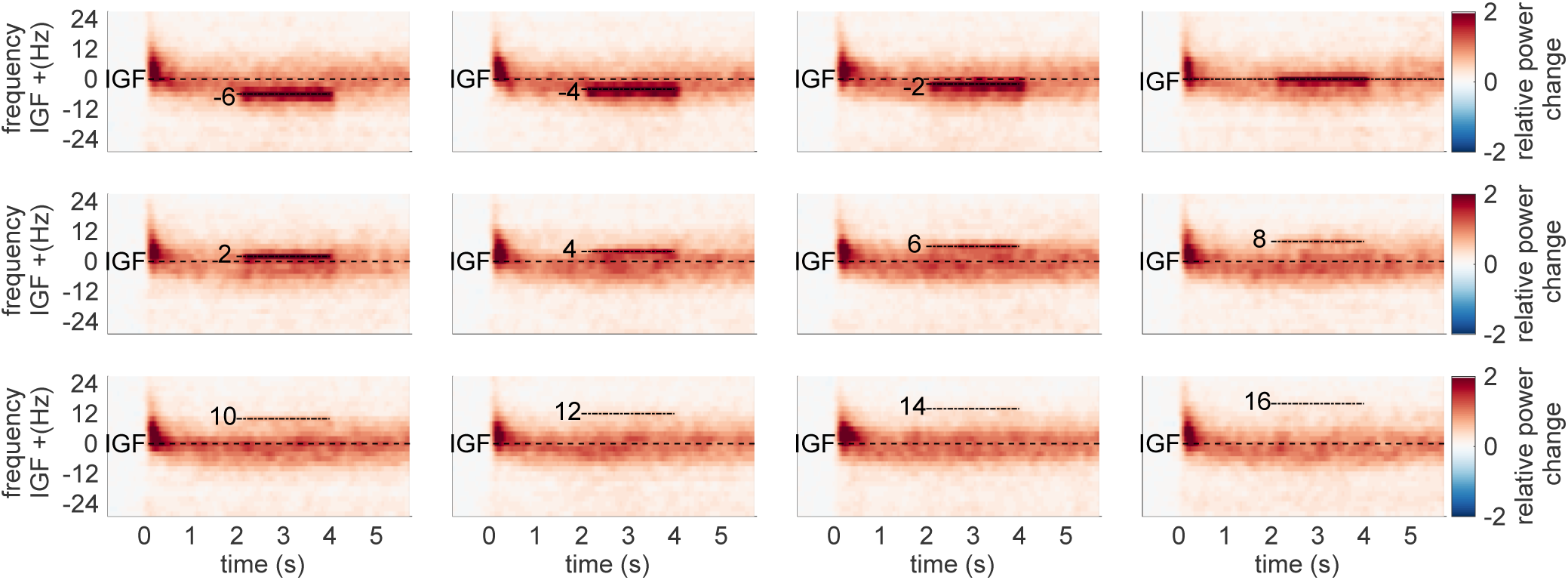
Grand-average TFRs of power after aligning to the IGF for each subject in the *flicker&gratings* condition. The stimulation frequencies (from -6 to 16 Hz relative to the IGF) are indicated by dashed horizontal lines. As suggested by the single subject TFRs in Figure 6, the endogenous gamma oscillations and the flicker response seem to be coexistent. Thus, there is no obvious indication of the photic drive being able to capture the dynamics of the gamma oscillations.

In addition to the narrow-band gamma oscillations, the gratings elicited a rhythmic response at 4 Hz, i.e. the velocity of the concentric drift (not shown). Apart from that, we did not find any evidence for an intermodulation between the frequency of the movement and the photic drive.

### 3.5 Frequency analyses with a longer time window confirm robustness of the reported results

To assess the robustness of our results, we repeated the frequency analyses in the *flicker* and *flicker&gratings* condition with a 2s sliding time window. The longer window substantially increased the signal-to-noise ratio of the flicker response, to up to over 400% relative power change in the *flicker* condition and more than 600% in the *flicker&gratings* condition (not shown). Besides that, the analyses replicated our reported main finding: a reduction in response magnitude (power) with increasing frequency, in both conditions, following the same trend as depicted in Figure 5C and D. The 2 s sliding time-window did however not optimally capture the gamma power, which has a broader peak than the response to the photic drive. The 500 ms sliding window used in our reported analyses is therefore a good compromise, allowing both a reliable identification of a gamma peak frequency and a sufficiently high signal-to-noise ratio and frequency resolution of the flicker response (see Figure 6A).

### 3.6 Oscillatory gamma dynamics cannot be captured by frequency entrainment

Synchronisation of neuronal oscillations by rhythmic stimulation could be conceptualized as the entrainment of a self-sustained oscillator by an external force (e.g. Notbohm et al., 2016; Helfrich et al., 2019). Frequency entrainment is reflected by a change in frequency of the ongoing oscillations towards the rhythm of the drive. Visual inspection of the TFRs of power in Figure 6 and 7 do no indicate any modulation of the peak frequency of the gamma oscillations by the flicker response, suggesting that they do not synchronize. To quantify these observations, we investigated the power of the gamma oscillations before and during the photic drive (Figure 8) in the *flicker&gratings* condition. A central assumption of oscillatory entrainment is the existence of a ’synchronization region’ in the frequency range around the endogenous frequency of the oscillator, the so-called Arnold tongue (e.g. Pikovsky et al., 2003). Driving frequencies falling inside this synchronization region, will be able to modulate the dynamics of the self-sustained oscillator (also see Hutt et al., 2018). With this in mind, the following analyses only included flicker frequencies in the vicinity of the IGF. For each participant, we considered the relative power change induced by the moving gratings in the 0.5 - 1.5 s interval (T1) before the flicker onset and in the 2.5 - 3.5 s interval (T2) in which both the moving gratings and the photic drive were present. We investigated this for stimulation frequencies below the IGF (averaged power for -6 and -4 Hz) and above (averaged power for +4 and +6 Hz). Assuming a symmetric Arnold tongue centered at the IGF, as shown for entrainment in the alpha-band (Notbohm et al., 2016), we expected a reduction in power at the IGF in interval T2 compared to interval T1 for both higher and lower driving frequencies, i.e. an effect of time, but not frequency. Figure 8 depicts power change at the IGF for the factors stimulation frequency (drive*<*IGF and drive*>*IGF) and time interval (T1 and T2), averaged over the SOIs for each subject. In accordance with the TFRs in Figure 7, there is no meaningful indication for gamma power being reduced during the T2 interval as compared to the T1 interval, affirming the coexistence of the two responses. A factorial repeated-measures ANOVA did not reveal any significant main effects of the factors time (T1 vs T2) and frequency (drive*<*IGF vs drive*>*IGF), but a significant interaction effect (*F* (1, 21) = 5.09*, p* = 0.003*, η*^2^ = .003). These results were further investigated using a Bayesian repeated-measure ANOVA. The obtained Bayes factors (*BF*_10_) indicate that the variance in the data underlies the variability between participants, while the factor *time* (*BF*_10_ = 0.233) and both *time* and *frequency* (*BF*_10_ = 0.274) do not add any explanatory value. Evidence for the interaction effect *time:frequency* was found to be inconclusive (*BF*_10_ = 0.53), as was the main effect of frequency alone *BF*_10_ = 1.146). These results provide evidence against the expected reduction in gamma power during rhythmic photic stimulation at frequencies different from the IGF; suggesting that the flicker did not capture the oscillatory gamma dynamics.

**Figure 8:**
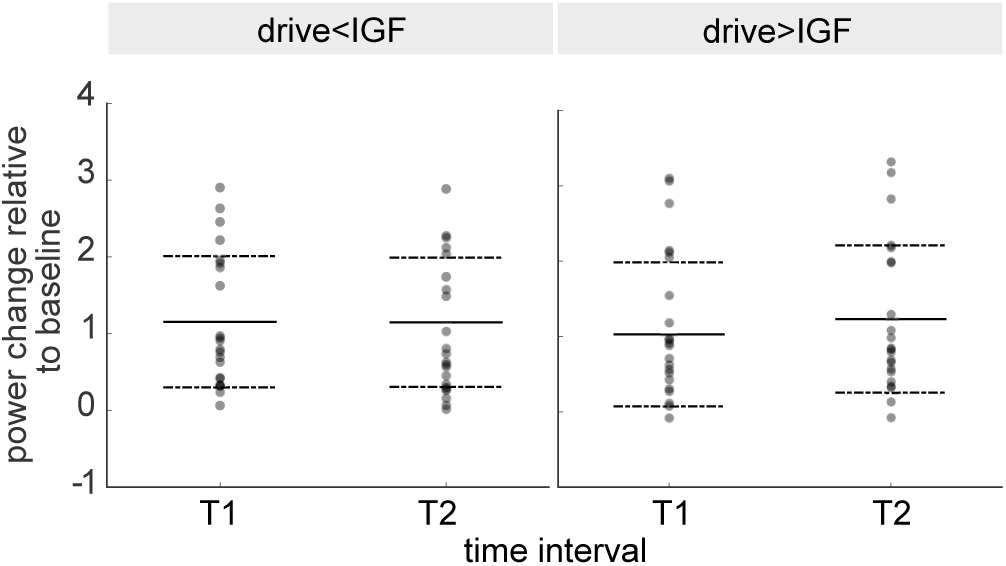
Power change relative to baseline at IGF in response to the moving grating stimuli before (T1; 0.5 - 1.5 s) and during application of the flicker (T2; 2.5 - 3.5 s), at frequencies below and above IGF (drive*<*IGF [-6, -4 Hz] and drive*>*IGF [+4, +6 Hz], respectively). Scatters demonstrate individual values, solid and dashed lines depict mean and standard deviation, respectively. The key finding is that power at T2 is not decreased compared to T1 for either of the frequency ranges, which is supported by a Bayesian repeated measures ANOVA (*BF*_10_ = 0.274).

### 3.7 Photic drive does not reliably modulate gamma phase

Synchronization of a self-sustained oscillator by an external force, can not only be described by a change in frequency, but also ’phase approximation’ or ’phase entrainment’ (Pikovsky et al., 2003). This phenomenon is reflected by a constant phase angle between the two oscillators over extended intervals, so-called *phase plateaus*. These might occur when the frequency of the driver is close to the endogenous frequency of the oscillator, i.e. within its Arnold Tongue (Tass et al., 1998; Pikovsky et al., 2003; Notbohm et al., 2016). When approaching the edge of the synchronization region, episodes of constant phase angles are interrupted by so-called *phase slips* that emerge when the self-sustained oscillator briefly unlocks from the driving force and oscillates at its own frequency. These phase slips will be observed as steps between the phase plateaus. The phase plateau analysis was implemented to complement the PLV analysis shown in Figure 5. The PLV quantifies the average synchrony between photodiode and neuromagnetic signal over trials using a 500 ms sliding time window. We hypothesized that in the case of oscillatory entrainment, the gamma oscillator in the *flicker&gratings* condition would alternate between locking on to the photic drive for a few cycles and slipping back to its endogenous rhythm. Due to the short duration of the gamma cycle (*∼*17.2 ms for a 58 Hz IGF), this intermittency would be smeared out by the sliding window. As there was no endogenous gamma oscillator in the flicker condition, such an intermittency was not expected. To investigate phase entrainment of the gamma oscillations by the photic drive, we inspected the phase angle between the photodiode and one, individually selected, occipital gradiometer of interest per participant. The time series of the phase were estimated per trial, separately for the two sensors, using a sliding time-window Fourier transform approach (ΔT = 3 cycles = 3/*f_flicker_* s; Hanning taper). Phase differences per trial were obtained by subtracting the unwrapped phase angle time series.

#### Phase angle between photodiode and MEG signal over time

Figure 9 illustrates the unwrapped phase angles between the MEG and photodiode signal during the photic drive at the IGF (here 58 Hz), in the *flicker* (A) and *flicker&gratings* condition (B), respectively, for the same representative participant shown in Figure 2A, 3A and 6. The colored line graphs depict individual trials. In both conditions, the MEG signal drifts apart from the photic drive, towards a maximum difference of 60 radians, i.e. a phase difference of about 9.5 cycles, by the end of the trial (A and B, top panel). Interestingly, the direction of the phase angle appears to change during some of the trials, suggesting spectral instability of the gamma oscillations. Furthermore, the graphs demonstrate a substantial inter-trial variability. This diffusion between trials, quantified for each participant as the standard deviation over trials at the end of the photic stimulation (t=2 in *flicker* and t=4 in *flicker&gratings* condition), converted from radiant to ms, is juxtapositioned in Figure 9C for the two conditions. It can be readily seen that the phase angles between the stimulation and MEG signal fan out highly similarly in absence and presence of the endogenous gamma oscillations.

**Figure 9:**
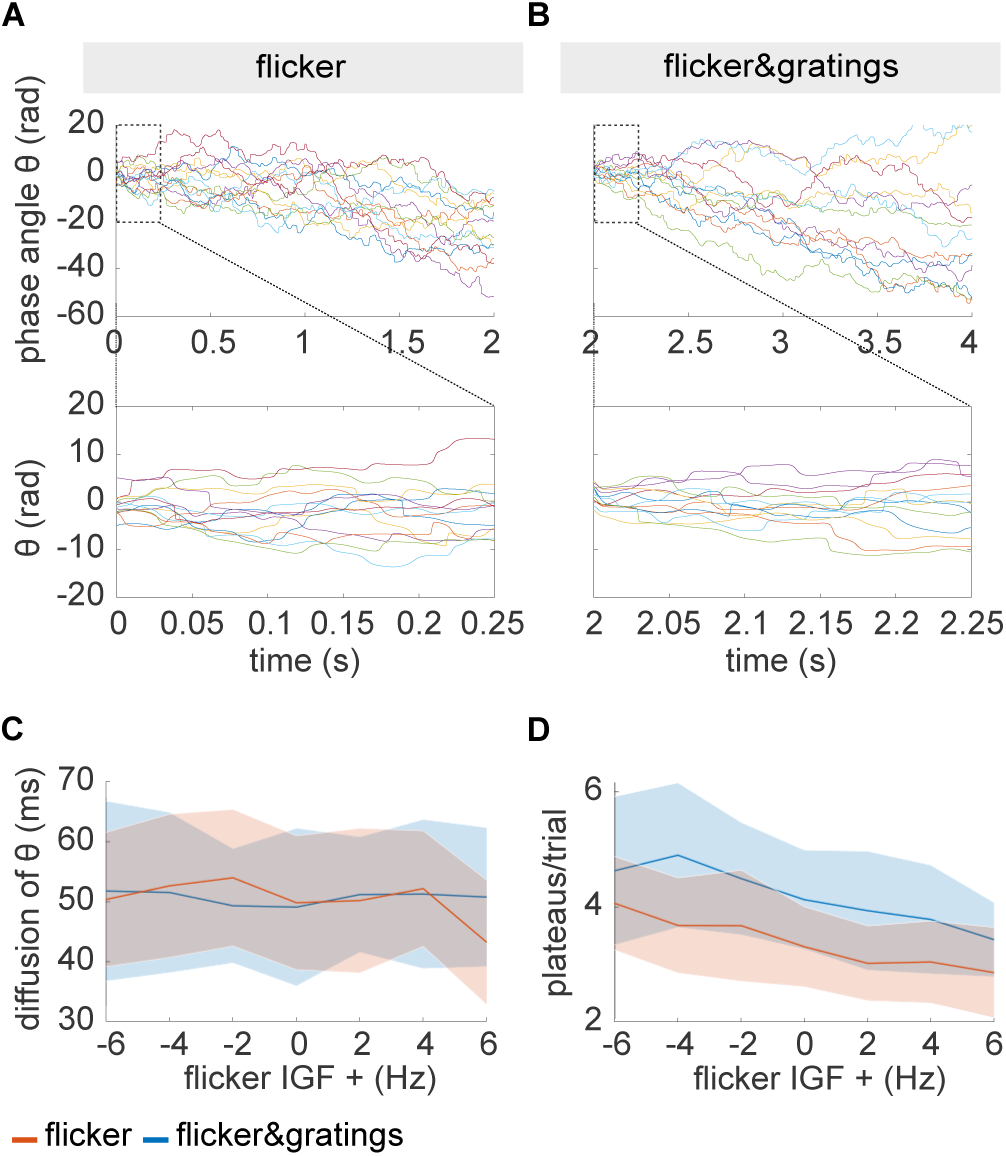
**A,B** Phase angle between photodiode and the MEG signal (one gradiometer of interest) at the IGF, for one representative participant; colored lines depict individual trials. **A** Phase angle *θ* in the *flicker* condition over duration of the flicker presentation (upper panel) and the first 250 ms (lower panel). The MEG signal drifts apart from the stimulation and can reach a maximum accumulated phase difference of 60 rad, i.e. 9.54 cycles, at the end of the stimulation and up to 15 rad, i.e. 2.39 cycles, in 250 ms. **B** The increase in phase difference over the time of the stimulation for the *flicker&gratings* condition (upper panel) and in the first 250 ms (lower panel). The diffusion of the phase difference across trials is similar to the *flicker* condition. Moreover, there is no clear difference in the number and length of phase plateaus between conditions, implying that the presence of the gamma oscillations does not facilitate entrainment at the IGF. **C** Fanning out across trials as a function of frequency aligned to IGF. Trials diffuse to a highly similar extent in both conditions and across frequencies. **D** Number of plateaus per trial as a function of frequency. While the *flicker&gratings* conditions exhibits more plateaus for all flicker frequencies, there is no indication that stimulation at the IGF results in comparably strong synchronization.

#### Phase plateaus

Visual inspection of the first 0.25 s of the phase angle times series, depicted in Figure 9A,B lower panel, does not suggest a relatively high number of phase plateaus in the *flicker&gratings* compared to the *flicker* condition, that would have been expected if the photic drive was able to entrain the endogenous gamma oscillator. Importantly, the graphs demonstrate the phase angles to reach values of over 2*π*, i.e. more than one cycle, within the duration of the first gamma cycle (17.2 ms), suggesting that even stimulation at the endogenous frequency of the oscillator cannot capture the gamma dynamics. To verify these observations for the entire sample, plateaus during stimulation at the IGF were identified based on the mean absolute gradient (⩽0.01 rad/ms, see equation 3) over the duration of one cycle of stimulation, i.e. 18 consecutive samples for a flicker frequency of 58 Hz. Figure 9D shows the average number of plateaus per trial as a function of flicker frequency aligned to IGF, averaged over participants. The shaded areas indicate the standard deviation. While the *flicker&gratings* condition exhibits more phase plateaus than *flicker* for all stimulation frequencies, the number of plateaus decreases similarly in both conditions with increasing frequency. Importantly, stimulation at the IGF did not result in the highest number of plateaus in either condition. these results are in line with the reported frequency analyses: responses to the photic drive in *flicker&gratings* show strong similarity to the *flicker* condition despite the presence of the gamma oscillator. The results affirm the observations presented in Figure 5A and B.

### 3.8 The sources of the gamma oscillations and the flicker responses peak at different locations

The coexistence of the endogenous gamma oscillations and flicker response suggest that these two signals are generated by different neuronal populations; possibly in different regions. To test this assumption we localized the respective sources using Linearly Constrained Minimum Variance spatial filters (LCMV; Veen et al., 1992). The covariance matrix for the spatial filters was estimated based on the -0.75 to -0.25 s baseline in both conditions, the 0.75 to 1.25 s interval with the moving gratings in *flicker&gratings* and the invisible flicker in the *flicker* condition, as well as the 2.75 to 3.25 interval in the *flicker&gratings* condition in which the flicker was applied to the grating stimulus. Note that for each participant, one common filter was used for source estimation in both conditions. Power values at the IGF and flicker frequencies, averaged up to 78 Hz, respectively for the *flicker&gratings* and *flicker* condition, were estimated based on the Fourier Transform. To extract power at the IGF and flicker frequencies, power change was computed relative to the baseline interval at each of the 37,163 grid points using equation 1. To isolate the flicker response on the *flicker&gratings* condition, the flicker&gratings interval was contrasted to the moving grating interval. Figure 10 illustrates the grandaverage of the source localization for the gamma oscillations (A), the invisible flicker response (B) and the response to the flickering gratings (C). Consistent with previous work, the responses originate from mid-occipital regions (Hoogenboom et al., 2006; Zhigalov et al., 2019). It is worth noting that the sources of the gamma oscillations and response to the invisible flicker are relatively focal, while the activity induced by the flickering gratings extends more broadly over visual cortex. Using the MNI to Talaraich mapping online tool by Biomag Suite Web (MNI2TAL Tool) (see Lacadie et al., 2007, 2008), the peak of the gamma oscillations was located in the ventral part of the secondary visual cortex (V2, Brodmann area 18; MNI coordinates = [-6mm -100mm -8mm], grandaverage). The peak sources of the flicker responses in both conditions were found in the Calcarine Fissure, at a 2mm distance to the border of the primary (V1) and secondary visual cortex (in dorsal direction), suggesting that they are generated by neighboring, coherent sources in both hemispheres in and close to V1 (Belardinelli et al., 2012) (MNI coordinates: flicker [6mm -96mm 12mm]; flicker&gratings [6mm -100mm 0mm]). To compare the peak locations between the sources in a lower dimensional space, the identified 3D coordinates were projected along their first Principal Component (Herrmann et al., 2011). Dependent sample t-tests revealed a significant difference in location between the peak sources of the IGF and the invisible flicker responses *t*(21) = *−*3.091*, p* = 0.017, Cohen’s *d* = *−*0.845, 95% CI [*−*1.5 *−* 0.2]*, B*_10_ = 8.2, as well as to the flickering gratings relative to gratings, *t*(21) = *−*2.633*, p* = 0.023, Cohen’s *d* = *−*0.495, 95% CI [*−*0.89 *−* 0.09]*, B*_10_ = 3.45; with the Bayes Factors B_10_ revealing moderate evidence for the H_1_ (Quintana and Williams, 2018). There was no significant difference in location between the sources of the flicker responses in both conditions, *t*(21) = 0.732*, p* = 0.472*, B*_10_ = 0.28, with the Bayes Factor providing moderate evidence for the H_0_. Note that all t-values were Benjamini-Hochberg-corrected for multiple comparisons. In light of the coexistence of the two responses observed in Figure 6 and 7, these results support the notion that gamma oscillations and flicker responses are generated by different neuronal populations.

**Figure 10:**
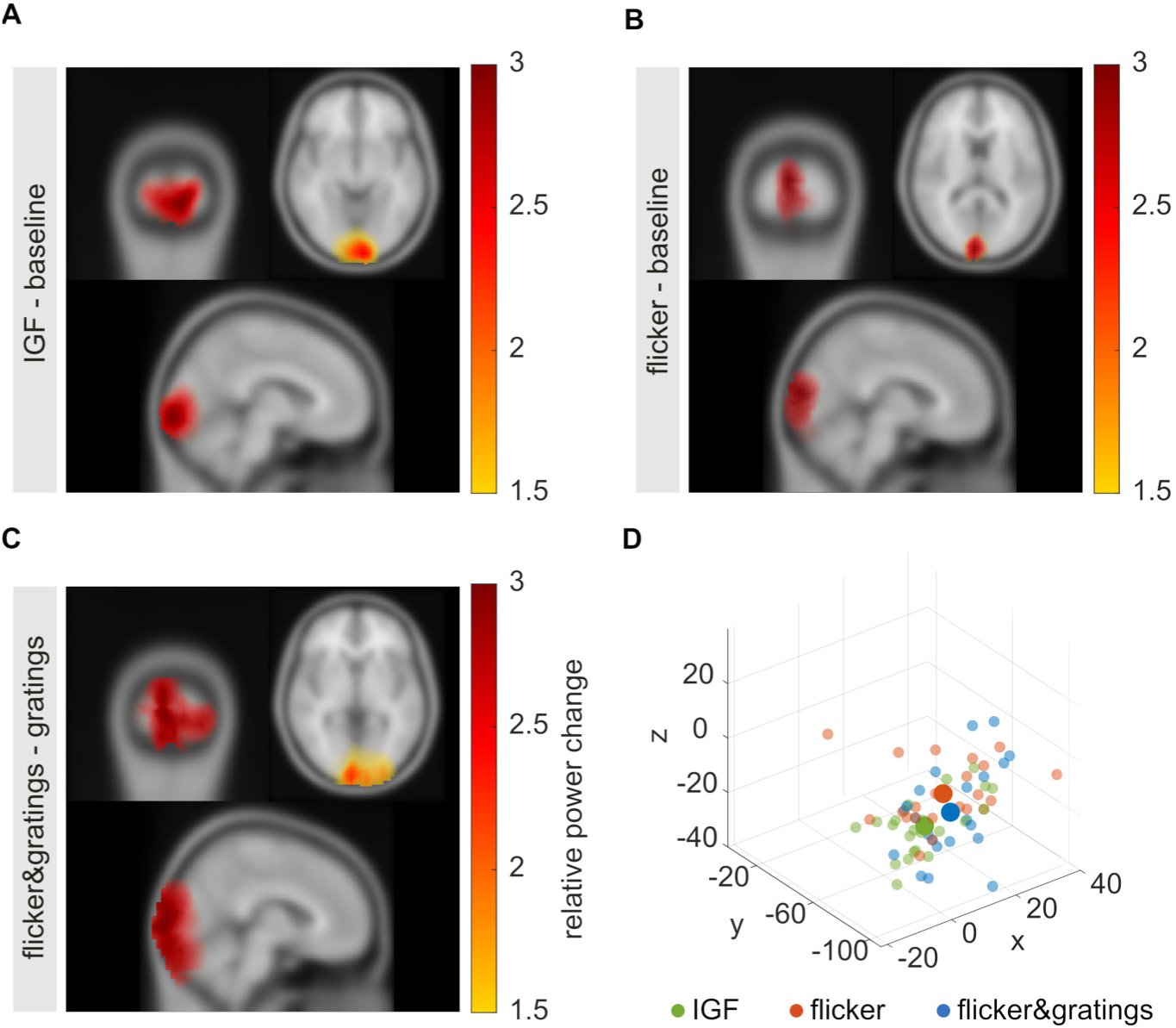
Source estimates using the LCMV beamformer approach mapped on a standardized MNI brain. **A** Source estimation of the visually induced gamma oscillations (power change relative to baseline), with the peak of the source identified at MNI coordinates [-6mm -100mm -8mm]. **B** Source estimation of the flicker response (relative to baseline), with the average peak source at [6mm -96mm 12mm] (in Calcarine Fissure). **C** Source estimation of the flicker response in the *flicker&gratings* condition (relative to the gratings interval), with the average peak source at [6mm -100mm 0mm] (in Calcarine Fissure). **D** Coordinates of the identified peak sources for all participants (small scatters) and grandaverage (large scatters) for the IGF, and the flicker responses in the *flicker* and *flicker&gratings* condition (green, orange and blue, respectively). The peak sources of the flicker responses are adjacent, while the gamma sources tend to peak at inferior locations.

## 4 Discussion

In this MEG study, we explored resonance and entrainment in the human visual system in response to a rapid photic drive *>*50 Hz. Strong, sustained gamma oscillations were induced using moving grating stimuli (Hoogenboom et al., 2006, 2010; Van Pelt and Fries, 2013; Muthukumaraswamy and Singh, 2013) and used to identify each participant’s gamma frequency. The superposition of the flicker and the gratings allowed us to investigate whether the flicker could entrain endogenous gamma oscillations. The photic drive induced responses for frequencies up to *∼*80 Hz, both in presence and absence of grating-induced endogenous gamma oscillations. To our surprise, we did not find evidence for resonance, i.e. an amplification of an individually preferred frequency in the range of the rhythmic stimulation, in either condition, despite the IGF being above 50 Hz in all participants. Moreover, there was no indication that the endogenous gamma oscillations synchronized with the rhythmic stimulation, i.e. no evidence for entrainment. Despite their differences, brain activity in the two conditions show strong similarities in the phase and frequency measures, supporting the notion that the flicker response coexists with the grating-induced oscillations. In accordance with these results, source estimation using Linearly Constrained Minimum Variance (LCMV) spatial filters (Veen et al., 1992), suggests that the neuronal sources of the flicker responses in both conditions and the endogenous gamma oscillations peak at different locations in visual cortex.

### 4.1 Flicker responses do not entrain the gamma oscillator

While the sources of the gamma oscillations and the response to the (nearly) invisible flicker did overlap in occipital cortex, their peak coordinates were found to be significantly different. Relative power change at the IGF peaked at sources inferior to the flicker responses in both conditions, and was located in the left secondary visual cortex (V2) using the MNI2TAL online tool (see Lacadie et al., 2007, 2008). The flicker peak sources were located in the Calcarine Fissure, in close proximity to the primary visual cortex (V1). These results are in line with the coexistence of the endogenous oscillations indicated by the time-frequency analyses and might be the result of the filter properties of synaptic transmission as the flicker response propagates in the visual system (see Kuffler, 1953; Hawken et al., 1996; Carandini et al., 1997; Ringach, 2004; Cormack, 2005; Shadlen and Movshon, 1999). Low-pass filtering at the transition from the thalamus to V1 (Connelly et al., 2015) might attenuate the photic drive at frequencies above 80 Hz, leading to an absence of measurable responses in this range. Low-pass filter properties in V1 in projections from granular layers (L4a, 4c*α* and 4c*β*) to supragranular (L2/3, 4b) and infragranular layers (L5,6) (Hawken et al., 1996; Douglas and Martin, 2004; Fröhlich, 2016) might have prevented the flicker response to converge to the neuronal circuits generating the endogenous gamma rhythms. This idea is supported by intracranial recordings in macaques showing the strongest gamma synchronization in response to drifting grating stimuli in V1 in supragranular layers (L2/3 and 4B) (Xing et al., 2012), whereas steady-state responses to a 60 Hz photic flicker were localised in granular layer 4c*α* (Williams et al., 2004). While plausible, these interpretations are conjectural based on the present data. Recent findings by Drijvers et al. (2020), providing evidence for non-linear integration of visual and auditory rapid frequency tagging signals in frontal and temporal regions, challenge the notion that the flicker response might not propagate beyond V1. Pairing the current paradigm with intracranial recordings in non-human primates would allow to test the filtering properties without the limitations imposed by the inverse problem in the source localization of neuromagnetic signals (Baillet, 2013).

#### Flicker responses might not be wired to inhibitory interneurons orchestrating the endogenous gamma rhythm

Computational models, as the one demonstrated by Tiesinga (2012, also see Lee and Jones, 2013), would be suitable to investigate whether the grating-induced gamma oscillations and flicker response are likely to be generated by neuronal circuits whose wiring is not conducive to entrainment. As the properties of neuronal gamma oscillations have been repeatedly shown to depend on rhythmic inhibition imposed by inhibitory interneurons (e.g. Wilson and Cowan, 1972; Bartos et al., 2007; Buzśaki and Wang, 2012; Lozano-Soldevilla et al., 2014; Kujala et al., 2015), entrainment should only be achieved when the flicker response is able to modulate their activity. Indeed, Cardin et al. (2009) show resonance in the gamma range to optogenetic stimulation of fast-spiking interneurons, but not to stimulation of pyramidal cells (also see Tiesinga, 2012). We therefore suggest that the photic stimulation applied in our study drives the pyramidal cells in early visual cortex. As in the optogenetic study by Cardin et al. (2009), this drive is not sufficiently strong to entrain the GABAergic interneurons. This interpretation is contrasted to the findings of Adaikkan et al. (2019) who demonstrate that a non-invasive 40 Hz flicker evokes neuronal processes counteracting neuro-degeneration (Singer et al., 2018; Adaikkan et al., 2019). However, it should be noted that the authors understand entrainment as the neural response to rhythmic stimulation, rather than a synchronization of ongoing oscillations to an external drive (Adaikkan and Tsai, 2020). While our findings do not question the authors’ compelling evidence that fast photic stimulation impacts neurocircuits and glia, the current study shows that it is not trivial to attribute these effects to entrainment of endogenous gamma oscillations.

### 4.2 Coexistence of flicker responses and oscillations versus oscillatory entrainment

The current study was inspired by studies reporting that a visual flicker in the alpha-band can capture the oscillatory dynamics of the visual system: resonance at distinct frequencies (Herrmann, 2001; Schwab et al., 2006; Gulbinaite et al., 2019, see Rager and Singer 1998 for flicker responses in cat visual cortex), amplitude and phase effects outlasting the stimulation interval (Spaak et al., 2014; Otero et al., 2020) and an “Arnold Tongue” relationship between stimulation intensity, distance to the individual alpha frequency and flicker-response-synchrony (Notbohm et al., 2016). Unlike the works listed above, we did not find any indication for a synchronization or resonance of endogenous oscillations in the gamma band to the visual stimulation. Recent studies applying photic stimulation in the alpha band, have pointed to a coexistence of endogenous alpha oscillations and flicker responses, similar to the one we report here for the gamma band. While retinotopic alpha modulation has been associated with suppression of unattended stimuli, allocating attention to a stimulus flickering in the alpha band results in enhanced, phase-locked activity (Keitel et al., 2019; Gundlach et al., 2020, also see Antonov et al. 2020; Friedl and Keil 2020 for stimulation at frequencies adjacent to the alpha-band). While the presented study does not allow nor aim to make generalized claims in favor or against neuronal entrainment, it is worth noting that the ability of rhythmic sensory stimulation to entrain endogenous oscillations is still a matter of debate.

### 4.3 Limitations & Generalizability

#### Interpretation of the different locations of the peak sources

The results of the LCMV beamforming are in line with the notion that gamma oscillations and flicker response are generated by sources at different locations. Yet, due to the ill-posed inverse problem (Baillet, 2013) and the merging of coherent sources when using the LCMV approach (Belardinelli et al., 2012) these source estimates should be interpreted with caution. Figure 10 illustrates that the sources of the flicker response in the *flicker&gratings* condition extended more broadly over visual cortex than the sources of the gamma oscillations and invisible flicker response, which might be the result of the flickering rings stimulating different receptive fields (Gur and Nodderly, 1997). While our results suggest a coexistence of the gamma oscillations and flicker response, we do not exclude they interact. These limitations do not seriously challenge our interpretation that the neuronal populations generating the flicker response do not entrain the activity of the neurons generating the endogenous gamma rhythm. Firstly, it is reasonable to assume that the peak sources reflect the flicker response, which tends to be stronger than the endogenous gamma oscillations (see Figure 6 and 7). Secondly, the significant difference between the peak locations of the gamma oscillations and flicker response in the *flicker&gratings* condition provides circumstantial evidence for the notion that the two responses emerge from different neuronal populations, despite being elicited by the same stimulus; albeit there is also overlap between the sources. Intracranial recordings in nonhuman primates or humans would be useful to substantiate this interpretation.

#### Strong flicker responses despite limited stimulation strength

The number of conditions that have been tested in this paradigm, i.e. 40 frequency*×*condition combinations, imposed limitations on the maximum number of trials per condition (N=15) and the duration of the stimulation (2 seconds). Stimulation strength was limited to a contrast of 66% peak to trough, ensuring equal luminance across conditions. Due to these limitations, one might be concerned that the absence of oscillatory entrainment was caused by the limited magnitude of the photic drive. However, we found the flicker to induce strong responses of up to 400% in the *flicker&gratings* condition and over 200% in the flicker condition (e.g. see Figures 4 and 5). In light of these response magnitudes, we argue that the absence of evidence for entrainment cannot be explained by the photic drive being too weak.

#### Generalizability of the current findings to gamma oscillations associated with visual perception

The use of drifting gratings is a standard approach to induce strong narrow-band gamma oscillations in humans (e.g. Hoogenboom et al., 2006, 2010; Muthukumaraswamy and Singh, 2013; Van Pelt et al., 2012; Van Pelt and Fries, 2013; Michalareas et al., 2016) and nonhuman primates (e.g. Womelsdorf et al., 2006; Bosman et al., 2012; Buffalo et al., 2011). One might argue that the conclusions presented here only apply to these stimuli and that entrainment could have been achieved using more complex stimuli such as natural images or faces. We find this very unlikely for the following reasons: Natural stimuli have been argued to induce gamma-band responses that are characterized by broadband activity (Ray and Maunsell, 2010; Hermes et al., 2015a,b, but also see Brunet et al. 2014; Bartoli et al. 2019; Brunet and Fries 2019). This is likely explained by the fact that gamma power and frequency depend on stimulus properties such as contrast, size and orientation (Schadow et al., 2007; Ray and Maunsell, 2010; Jia et al., 2013; Muthukumaraswamy and Singh, 2013). As these factors vary greatly within a natural image, the net result of the oscillatory activity in the gamma-band is a broadband response. Moving gratings have been shown to induce stronger gamma oscillations than their stationary counterparts (Muthukumaraswamy and Singh, 2013; Perry et al., 2013) and were therefore chosen for the current paradigm. We expected the flicker responses to be substantially stronger than the grating-induced gamma oscillations, which is confirmed by Figure 6 and 7. Had we relied on stationary gratings, the photic drive might have overshadowed weaker gamma-band activity. Moreover, the frequencies of the endogenous gamma rhythms have been found to be higher for moving than for stationary gratings (Muthukumaraswamy and Singh, 2013; Perry et al., 2013). As our study aimed to investigate entrainment by a flicker with minimal visibility, the IGFs had to be relatively high to be in the range of feasible stimulation frequencies. While the gratings’ concentric drift did induce a rhythmic response at 4 Hz, there was no evidence for an intermodulation with the flicker frequencies, nor an indication that the *flicker&gratings* condition was lacking spectral precision. Another concern might be that grating stimuli do not engage downstream regions to the same extent as complex stimuli; as such they might be generated in specialized neuronal circuits. However, a number of studies in both human and non-human primates have demonstrated that attended as well as unattended gratings induce gamma oscillations that propagate to downstream areas along the ventral (V4 and inferotemporal cortex) and dorsal stream (area V5 and V7) (Buffalo et al., 2011; Bosman et al., 2012; Bastos et al., 2015; Michalareas et al., 2016). For the reasons outlined above, we argue that moving grating stimuli created the optimal conditions to investigate gamma-band entrainment, as these induced strong, sustained, narrow-band gamma oscillations reflecting individual oscillatory dynamics (also see Hoogenboom et al., 2006; Van Pelt and Fries, 2013).

### 4.4 Concluding remarks

Our results suggest that rapid photic stimulation does not entrain endogenous gamma oscillations and can therefore not be used as a tool to probe the causal role of gamma oscillations in cognition and perception. However, the approach can be applied as Rapid Frequency Tagging (RFT) to track neuronal responses without interfering, for instance, to investigate covert spatial attention (Zhigalov et al., 2019), multisensory integration (Drijvers et al., 2020) and parafoveal reading (Pan et al., 2020).

## Acknowledgements

This study was funded by a James S. McDonnell Foundation Understanding Human Cognition Collaborative Award (grant number 220020448), the Wellcome Trust Investigator Award in Science (grant number 207550), a BBSRC grant (BB/R018723/1), as well as the Royal Society Wolfson Research Merit Award (awarded to O.J.). The authors are grateful to Prof Veikko Jousmaki for providing the light-to-voltage converter and to Jonathan L. Winter, Nina Salman, Ludwig Barbaro, and Roya Jalali for providing help with the MEG recordings and MRI scans. The authors further thank Dr Simon Hanslmayr, Dr Geoffrey Brookshire and Dr Florian Kasten for feedback on the project and manuscript.

